# A divergent nonsense-mediated decay machinery in *Plasmodium falciparum* is inefficient and non-essential

**DOI:** 10.1101/2021.04.14.439394

**Authors:** Emma McHugh, Michaela S. Bulloch, Steven Batinovic, Drishti K. Sarna, Stuart A. Ralph

## Abstract

Nonsense-mediated decay (NMD) is a conserved mRNA quality control process that eliminates transcripts bearing a premature termination codon. In addition to its role in removing erroneous transcripts, NMD is involved in post-transcriptional regulation of gene expression via programmed intron retention in metazoans. The apicomplexan parasite *Plasmodium falciparum* shows relatively high levels of intron retention, but it is unclear whether these variant transcripts are functional targets of NMD. In this study, we use CRISPR-Cas9 to disrupt and epitope-tag two core NMD components: *Pf*UPF1 (PF3D7_1005500) and *Pf*UPF2 (PF3D7_0925800). Using RNA-seq, we find that NMD in *P. falciparum* is highly derived and requires UPF2, but not UPF1 for transcript degradation. Furthermore, our work suggests that the majority of intron retention in *P. falciparum* has no functional role and that NMD is not required for parasite growth *ex vivo*. We localise both *Pf*UPF1 and *Pf*UPF2 to puncta within the parasite cytoplasm, which may represent processing bodies - ribonucleoparticles that are sites of cytoplasmic mRNA decay. Finally, we identify a number of mRNA-binding proteins that co-immunoprecipitate with the NMD core complex and propose a model for a divergent NMD that does not require *Pf*UPF1 and incorporates novel accessory proteins to elicit mRNA decay.

## INTRODUCTION

Nonsense-mediated decay (NMD) is a conserved mRNA degradation pathway that detects and eliminates transcripts containing a premature termination codon (PTC). Destabilisation of nonsense transcripts occurs in almost all studied eukaryotes. Yet despite conservation of the three core NMD proteins (UPF1, UPF2 and UPF3b) in all major eukaryotic groups, the mechanism of NMD, especially outside of opisthokonts, is unclear. Studying NMD in diverse organisms is key to understanding how this ancient quality control mechanism has evolved.

During translation, ribosome stalling at a PTC can trigger NMD. There are two general models that describe how a termination codon is identified as premature and cause the initiation of NMD. These models are termed exon junction complex (EJC)-dependent NMD, which predominates in mammals, and faux 3’-UTR NMD. The former relies on EJC proteins, which are deposited 20-24 nucleotides upstream of exon-exon junctions after splicing. In mammals, EJC-dependent NMD occurs when an EJC is present > 50-55 nucleotides downstream of a PTC during translation termination (Fig. 1A) (1, 2). Faux 3’-UTR NMD is observed in animals such as *Drosophila melanogaster* and *Caenorhabditis elegans* which have EJCs but do not require them for NMD, and also in the yeast *Saccharomyces cerevisiae*, which lacks EJC components. This model of NMD posits that after encountering a PTC, the absence of a nearby normal 3’UTR, poly(A) tail and poly(A)-binding protein destabilises mRNA (3). Other steps in NMD are less easily generalised by existing models due to low conservation of proteins between eukaryotic groups. These poorly understood steps include the activation of the RNA helicase UPF1 by phosphorylation (and whether this is important for NMD in all species), and the degradation of PTC-containing transcripts by nucleases.

**Figure 1.**
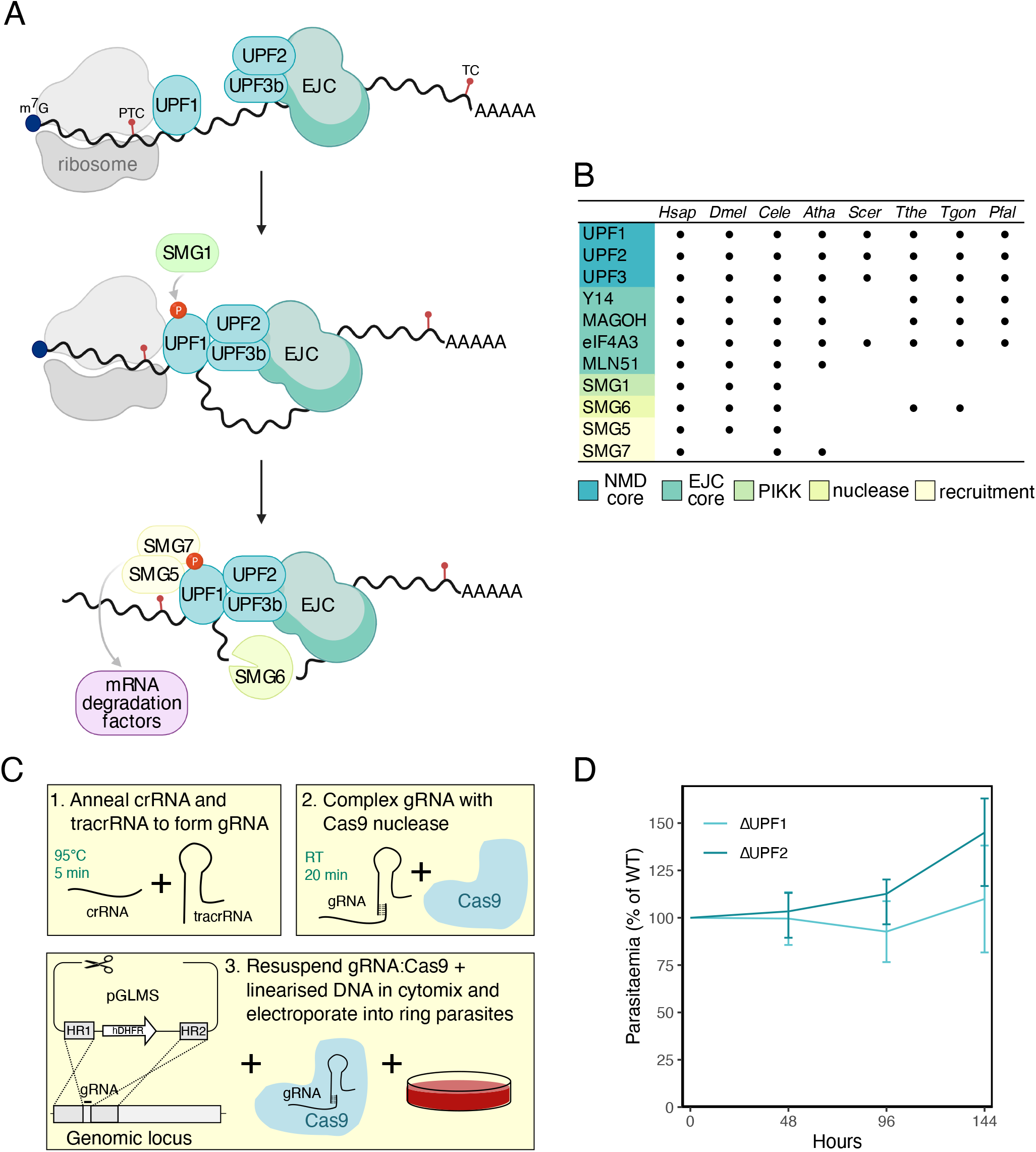
**(A)** Model of EJC-dependent NMD in humans. After ribosome stalling at a PTC, UPF2 and UPF3b associated with the downstream (>50-55nt) EJC will bind to UPF1. SMG1 phosphorylates and activates UPF1, resulting the recruitment of SMG5 and SMG7 (involved in the further recruitment of mRNA decay factors) and the endonuclease SMG6. **(B)** Presence of NMD-associated proteins in selected species (*H. sapiens, D. melanogaster, C. elegans, A. thaliana, S. cerevisiae, T. thermophila, T. gondii* and *P. falciparum*). Proteins are grouped based on their role in NMD as determined in mammals. The NMD core are required for initiation of NMD. EJC core proteins are deposited upstream of exon-exon boundaries after splicing and are required for EJC-dependent NMD. SMG1 is a phosphatidylinositol 3-kinase-related kinase (PIKK) which phosphorylates UPF1. SMG6 is an endonuclease. Both SMG5 and SMG7 are involved in recruitment of UPF1 to cytoplasmic RNA degradation sites such as the exosome. This table was compiled using genes identified in (26). **(C)** Schematic protocol for disruption of the *Pf*UPF1 and *Pf*UPF2 genes using the Alt-R CRISPR Cas9 system (Integrated DNA Technologies). The gene-specific crRNA is annealed to the Cas9-binding tracrRNA to form gRNA (1) which is then complexed with recombinant Cas9 to form the gRNA:Cas9 RNP (2). The gRNA:Cas9 RNP is then mixed with a linearised DNA repair template encoding two homology regions (HRs) flanking a human dihydrofolate reductase (hDHFR) expression cassette. Finally, the DNA + gRNA:Cas9 mixture is electroporated into ring-stage parasites (3). **(D)** Growth analysis of Δ*Pf*UPF1 and Δ*Pf*UPF2. Parasitaemia was measured by flow cytometry over 3 asexual replication cycles (144 h) and is plotted as percentage of WT parasitaemia. Error bars represent SEM (n = 3).

PTC-containing transcripts that are subject to NMD can be generated in a number of ways, including nonsense mutation, transcriptional error, or alternative splicing of pre-mRNA. Alternative splicing is widespread in eukaryotes and can refer to a number of different splicing events such as intron retention, exon skipping or alternative splice site usage. We use the term ‘alternative splicing’ to refer to the generation of both functional splice variants and mis-splicing/spliceosome errors. Alternative splicing frequently produces transcripts that contain a PTC and the degradation of these transcripts by NMD has been proposed as a method for the regulation of gene expression (4, 5). This process has been best characterised in the auto-regulation of splicing factors such as serine-arginine (SR) proteins (6). Programmed intron retention leading to NMD may also contribute to global regulation of mRNA abundance in humans, yeast, and plants, and has been reported to control processes such as differentiation, development and immune responses (7-9).

The malaria parasite *P. falciparum* has high levels of intron retention compared to other forms of alternative splicing, which is markedly different from the proportions of alternative splice variants in humans (10). We therefore investigated whether intron retention coupled to NMD has a role in the regulation of gene expression in *P. falciparum*. High levels of observed intron retention could also indicate a lack of NMD, so we set out to ascertain whether the core NMD protein homologues in *P. falciparum* have canonical functions. In this study, we perform CRISPR-Cas9 editing of *P. falciparum* using commercially available recombinant Cas9 and guide RNA, highlighting an efficient and cost-effective transfection strategy. We describe a highly derived form of NMD that appears to have only a minor impact on the steady state abundance of most transcripts, and which is not required for asexual parasite proliferation *ex vivo*. We propose a model for how PTC-containing transcripts are coupled to NMD surveillance machinery in *P. falciparum*.

## RESULTS AND DISCUSSION

### Efficient disruption of conserved NMD core proteins in *P. falciparum* using commercial CRISPR-Cas9 ribonucleoproteins

The core NMD proteins UPF1, UPF2 and UPF3 are present in diverse eukaryotes (Fig. 1B). Although the NMD core complex is highly conserved, there are some exceptions in excavates such as *Giardia* spp., in which spliceosomal introns are extremely rare (11). Outside of the three core NMD proteins, some other proteins known to be involved in metazoan NMD have been lost in other eukaryotic lineages. For example, the EJC core proteins are required for EJC-dependent NMD in mammals, however components of this complex have been lost in organisms that undertake EJC-independent NMD, such as *S. cerevisiae* and *Tetrahymena thermophila* (12, 13). Additionally, the SMG1 kinase that phosphorylates and activates metazoan UPF1 for NMD (14) has been lost in *Arabidopsis thaliana* (but not all plants), *S. cerevisiae* (although it is present in other fungal linages) and alveolates including *P. falciparum* (15). Despite this, global phospho-proteomics indicates that *Pf*UPF1 is phosphorylated (16) (although the kinase is unknown) and it is not clear whether this phosphorylation is important for *Pf*UPF1 activity in NMD.

To our knowledge, NMD has been studied in only two alveolates: the ciliates *T. thermophila* and *Paramecium tetraurelia*. We are interested in the parasitic phylum Apicomplexa, which encompasses parasites such as *Toxoplasma gondii* and the human malaria parasite *P. falciparum*. Previous studies have shown that *Plasmodium* spp. rely on post-transcriptional regulation for parasite development, including processes such as mRNA sequestration (17) and alternative splicing (18). Some studies have speculated that NMD coupled to intron retention - the predominant form of alternative splicing in apicomplexans - coupled to NMD could contribute to widespread regulation of mRNA abundance (10, 19) and hence we are interested in characterising NMD in *P. falciparum*.

We targeted two NMD genes for disruption: *Pf*UPF1 and *Pf*UPF2 (PlasmoDB IDs: PF3D7_1005500 and PF3D7_0925800). These genes were identified as the putative homologues of UPF1 and UPF2 in a bioinformatic study that catalogued RNA-binding proteins in *P. falciparum* (20). For CRISPR editing in *P. falciparum*, the Cas9 nuclease is usually encoded on a plasmid which is transfected in parallel with the homologous repair template and guide RNA expression template. Two or more of these features may be present on the same plasmid, and many variants of this strategy are currently in use with permutations of selectable markers, Cas9, guide RNA and repair templates in different arrangements (21-23). However, such plasmid construction can be complex and time consuming, and one widely-adopted method for insertion of the guide RNA requires expensive reagents and special DNA purification procedures for the restriction enzyme BtgZI (24). In other cell types and organisms an alternative to intracellular expression of the Cas9 and guide RNA is delivery of the Cas9:guide RNA complex, i.e. ribonucleoproteins (RNPs), for example by electroporation or direct injection. Ribonucleoprotein delivery has been used for CRISPR editing in model organisms such as *C. elegans, Arabidopsis thaliana*, zebrafish and mice (25-27). These RNP complexes can be delivered to human embryonic stem cells, T cells and other cell types by electroporation (28, 29).

We made use of the Alt-R® CRISPR-Cas9 system (Integrated DNA Technologies) to disrupt the coding region of *Pf*UPF1 and *Pf*UPF2 (Fig. 1C). Custom guide RNAs were synthesised as crRNA and annealed with the Cas9-binding tracrRNA, followed by complexing with recombinant *S. pyogenes* Cas9 nuclease (Integrated DNA Technologies). The resulting RNP was electroporated into *P. falciparum* with a linear DNA repair template. Integration was confirmed by PCR, demonstrating that electroporation of *P. falciparum* with commercially available ribonucleoproteins can lead to successful genome editing (Fig. S1). This method saved time on cloning of guide RNAs into unwieldy plasmids and also saved on costly plasmid assembly reagents and large-scale DNA preparation.

### NMD is not required f or parasite replication or maintenance of steady-state mRNA levels

In keeping with a genome-wide mutagenesis study (30), we found that both *Pf*UPF1 and *Pf*UPF2 were dispensable for asexual parasite growth after disruption by CRISPR/Cas9 (Fig. 1D). In humans, disruption of UPF1 leads to an accumulation of transcripts containing PTCs (31). These PTC-containing transcripts can arise from transcriptional or splicing errors that would be detected by UPF1 during translation and subsequently degraded through the process of NMD. However, alternative splicing of transcripts so that they contain a PTC (e.g., intron retention or inclusion of a ‘poison exon’) can be important for regulation of transcript abundance. For example, some SR splicing proteins have been shown to autoregulate their expression in animals and fungi via NMD by directing alternative splicing of their cognate transcript (6, 31). We reasoned that if NMD occurs in *P. falciparum*, we would observe upregulation of transcripts modulated by alternative splicing coupled to NMD upon *Pf*UPF1 or *Pf*UPF2 disruption. To test this, we extracted RNA from WT, Δ*Pf*UPF1 and Δ*Pf*UPF2 parasites (three biological replicates) and performed Illumina mRNA sequencing. We then tested for differential gene expression using the limma-voom method. Compared to WT, there were 15 and 13 differentially expressed transcripts in Δ*Pf*UPF1 and Δ*Pf*UPF2 respectively, defined as an adjusted p-value < 0.05 calculated using the ‘treat’ method with a required log-fold change > 1 (Table S1 and Table S2). Aside from detecting downregulation of *Pf*UPF1 in the Δ*Pf*UPF1 parasites as expected (Fig. 2A), variant antigens such as *Pf*EMP1 (*P. falciparum* erythrocyte membrane protein 1), RIFINs (repetitive interspersed family) and STEVORs (subtelomeric variable open reading frame) were also differentially expressed in both Δ*Pf*UPF1 and Δ*Pf*UPF2 (Fig. 2B) parasites. These variant antigens are members of multicopy gene families that are subject to stochastic transcriptional switching (32), and so are unlikely to represent changes due to knockout of NMD genes. We also assessed intron retention in Δ*Pf*UPF1 and Δ*Pf*UPF2 parasites using the ASpli R package and found that there was no differential usage of any introns (Table S3 and Table S4). Together, the lack of upregulation of specific transcripts or of particular introns suggests that NMD does not participate in targeted regulation of transcript abundance and nor does it produce a significant change in processing for any individual gene in asexual blood stages of *P. falciparum*.

**Figure 2.**
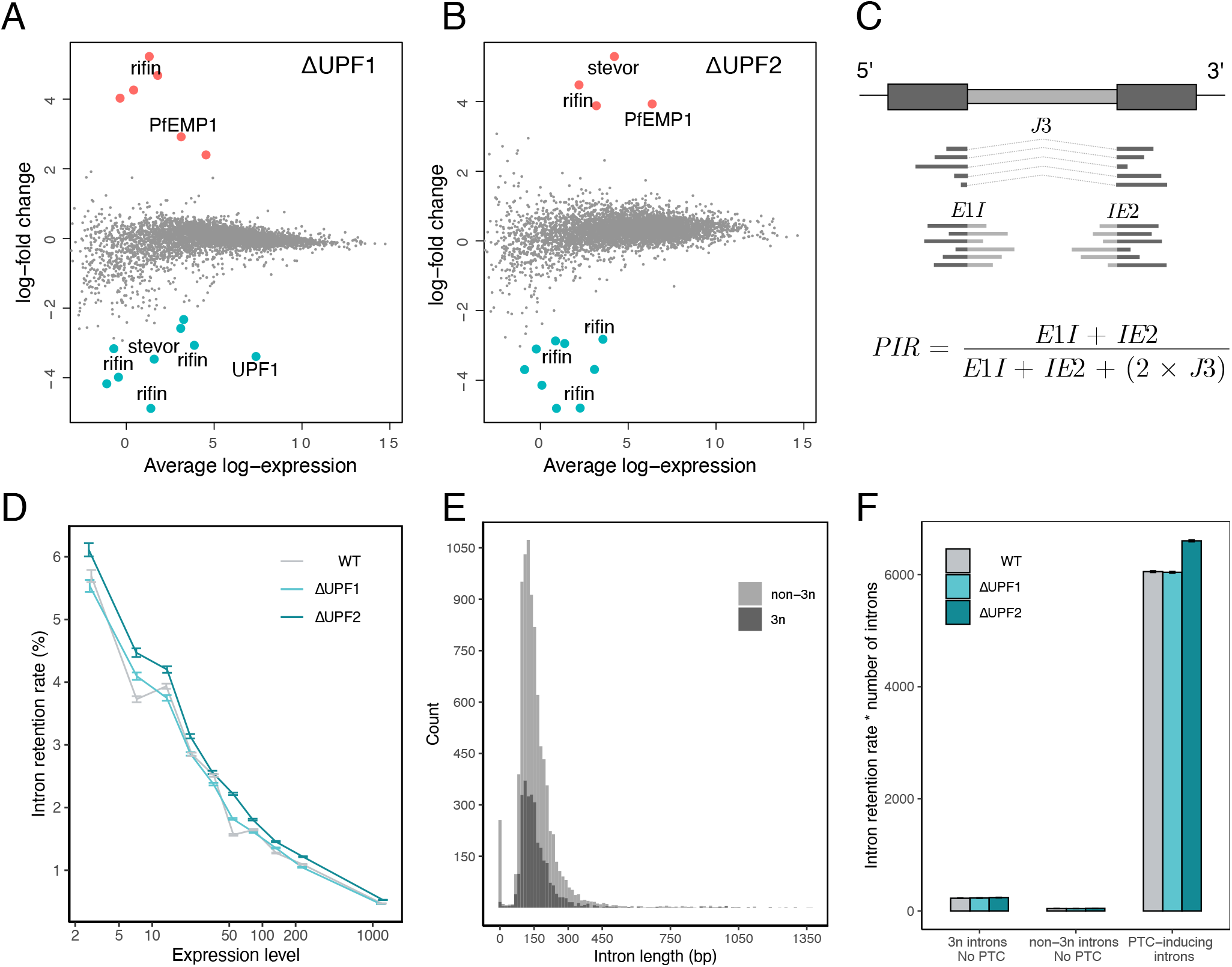
**(A + B)** Mean-difference plots showing differential gene expression in Δ*Pf*UPF1 and Δ*Pf*UPF2 parasites. Highlighted genes are considered significantly upregulated (red) or downregulated (blue) with an adjusted p-value < 0.05 (Benjamini-Hochberg) as calculated using the limma ‘treat’ method (log-fold-change > 1) with voom normalisation. **(C)** Schematic depicting the calculation of the global proportion of intron retention (PIR). An example gene with two exons (dark grey) and one intron (light grey) is shown. *J*3 reads span the spliced junction, *E*1*I* reads span the 5’ exon-intron boundary and *IE*2 reads span the 3’ intron-exon boundary. For a given expression level bin, reads were summed according to their designation as *E*1*I, J*3 or *IE*2, and the global PIR was then calculated using the formula shown. Figure adapted from the ASpli reference manual (April 27, 2020 release). **(D)** Introns (n = 7523) were grouped into 10 equal-sized bins based on expression level (FPKM) and the PIR was computed globally within each bin for WT, Δ*Pf*UPF1 and Δ*Pf*UPF2. Error bars represent 95% confidence interval. **(E)** Introns were classified into two groups: non-3n if the intron length is not a multiple of 3 (i.e. retention of the intron causes a frameshift) and 3n if the intron length is a multiple of 3. Introns were divided equally into 100 bins based on length (bp) before plotting. **(F)** Introns were classified into three groups based on the effect of retention: those that neither induce a frameshift nor PTC (3n introns = 143), introns that induce a frameshift but no PTC (non-3n introns = 34) and PTC-inducing introns (6725 introns). Intron retention was then calculated globally within each bin. For plotting, the intron retention rates and confidence intervals were multiplied by the number of introns in each of the three groups. Error bars represent 95% confidence interval.

### NMD degrades PTC-containing transcripts in *P. falciparum*

One function of NMD is quality control, i.e., identifying and inducing degradation of PTC-containing mRNAs that arise from transcriptional error, nonsense mutation, and mis-splicing. This ‘house-keeping’ role of NMD is conserved in diverse eukaryotes. It has also been suggested that the programmed generation of PTC-containing mRNAs via intron retention, and subsequent degradation through NMD, is an important method for global post-transcriptional regulation of expression (7, 9, 33). However, a study that examined intron retention from the perspective of the fitness cost of mis-splicing suggested that the vast majority of alternative splicing, including intron retention, is stochastic (34). This work, which was performed on datasets from *P. tetraurelia* and humans, showed that in general, intron retention is inversely correlated with expression level (34). This suggests that one of the main determinants of intron retention is selection for splice site strength, as it would be disadvantageous for highly expressed genes to consume cellular resources b y frequently producing aberrant transcripts. This would also imply that the majority of intron-containing mRNAs represent splicing errors, rather than a complex program of post-transcriptional regulation as has been suggested previously (7, 34). Using this framework, we investigated the relationship between intron retention rate and expression level in WT, Δ*Pf*UPF1 and Δ*Pf*UPF2 parasites. All genes were equally divided into ten bins based on expression level (FPKM) and a global intron retention rate (PIR) was calculated for all introns within each bin as shown in Fig. 2C. If NMD degrades erroneous PTC-containing transcripts, the intron retention rate should increase within a given bin when the key NMD factors are disrupted. Surprisingly, we found that this does not occur in Δ*Pf*UPF1 parasites - the intron retention rate 95% confidence interval (CI) is lower or overlaps the WT rate 95% CI in seven out of ten bins. This suggests that disruption of *Pf*UPF1 does not impair NMD. In contrast, the Δ*Pf*UPF2 intron retention rate 95% CI was higher than WT in for all except one bin, demonstrating a modest global increase in intron retention likely resulting from impaired NMD (Fig. 2D). Consistent with observations in *P. tetraurelia* and humans (34), we see an inverse correlation between intron retention and expression, arguing against widespread regulation of expression via intron retention and NMD in *P. falciparum*.

If the fitness cost of splicing errors is a main determinant of intron retention rate, we might also expect that each intron within a CDS with many introns would have a higher rate of correct (i.e., canonical) splicing than a single-intron CDS. We examined this in Δ*Pf*UPF1, Δ*Pf*UPF2 and WT parasites. As before, genes were binned by expression level and each CDS was further sorted into three groups, depending on the number of introns it contains (1-2 introns, 3-5 introns, >5 introns). The intron retention rate was then computed globally (Fig. S2). We observed no obvious relationship between intron number and intron retention rate in the lower six expression level bins. However, in the NMD-disrupted Δ*Pf*UPF2 parasites (Fig. S2C), at higher expression levels, introns in CDSs with 1-5 introns (n = 4191) had a higher overall retention rate than those within CDSs with >5 introns (n = 3332). Because there are many more reads mapping to highly expressed CDSs, the 95% CI of these intron retention rates are much smaller.

Considering that disrupting *Pf*UPF2 increases intron retention, we wanted to assess whether this was specific to PTC-inducing introns. If it was, this would provide further evidence for the existence of a canonical NMD pathway in *P. falciparum*. We divided introns into two groups (Fig. 2E); introns that induce a frameshift when retained (non-3n, 5894 introns, median length = 139) and introns with length that is a multiple of three that do not disrupt the reading frame when retained (3n, 2880 introns, median length = 141). We then classified each intron as either PTC-inducing or not, assuming it was the only intron retained in a given CDS. There were relatively few introns that do not induce a PTC when retained (n = 177) compared to those that do (n = 6725). The intron retention rate was then calculated globally for each category, with non-PTC-inducing introns further divided into 3n and non-3n (Fig. 2F). There was no difference between the intron retention rate of WT and Δ*Pf*UPF1 parasites for PTC-inducing introns (z-test, p = 0.13). However, the intron retention rate for PTC-inducing introns in Δ*Pf*UPF2 parasites was significantly increased, consistent with its role in NMD.

We next considered explanations for the discrepancy between the effects of disrupting *Pf*UPF1 compared to *Pf*UPF2. Most existing models of NMD in opisthokonts posit that initial recognition of premature termination by UPF1 is a crucial first step, followed by binding of UPF2, UPF3B and other factors. Under this model, disruption of *Pf*UPF2 should impair NMD (as we observed), but we would only expect that to be the case if disruption of *Pf*UPF1 also inhibited NMD. However, there are precedents for NMD without UPF1, such as a study in *Trypanosoma brucei* that found that although PTCs decreased reporter mRNA abundance (implying NMD), *Tb*UPF1 was not involved (35). Furthermore, a study in *S. cerevisiae* suggested that Upf2 could function in NMD independently from its Upf1-binding activity and that Upf2 may be able to function through its interactions with other RNA helicases (36).

We first re-confirmed that *Pf*UPF1 was genetically disrupted in Δ*Pf*UPF1 parasites by examining RNA-seq reads mapping to the UPF1 locus. In Δ*Pf*UPF1 parasites, no reads overlapped the Cas9 target site, indicating complete integration of the construct in the parasite population (Fig. 3A; chromosome 10, ∼0.2418 Mb). Reads mapping downstream of this site are likely due to spurious transcription initiation after the integrated drug cassette. The next possibility we considered is that the *P. falciparum* genome might contain another (as yet unannotated) UPF1. To search for such a gene, we performed a HMM search (http://hmmer.org/) using the *Homo sapiens* UPF1 sequence (UniProt ID: Q92900) as input and restricted results to the *Plasmodium* taxon. The second *P. falciparum* hit (after *Pf*UPF1) was PF3D7_0703500, which encodes a ∼234 kDa protein and is currently annotated in PlasmoDB as “erythrocyte membrane-associated antigen” (37). A mutagenesis study has suggested that PF3D7_0703500 is essential and it was identified with high-confidence as an mRNA-bound protein during the asexual blood stages (30, 38). Both *Pf*UPF1 and PF3D7_0703500 contain the conserved AAA ATPase domains (Pfam: AAA_11 and AAA_12) that are involved in the RNA helicase activity of UPF1 (Fig. S3). However, there is no obvious zinc binding UPF2-interacting domain in PF3D7_0703500. The protein sequence of PF3D7_0703500 is only 24.6% identical to the *H. sapiens* UPF1, whereas there is 46.1% identity between the protein which we refer to as *Pf*UPF1 and *H. sapiens* UPF1 sequences. Additionally, a phylogram with 13 protein sequences annotated as UPF1 (Fig. 3B, teal box), plus PF3D7_0703500 and three related proteins (Fig. 3B, pink box) shows that the PF3D7_0703500-related proteins form an outgroup distinct from the UPF1s. We therefore consider it possible but unlikely that PF3D7_0703500 protein is having a compensatory effect for canonical UPF1 function in the Δ*Pf*UPF1 parasites. We next performed a multiple sequence alignment with UPF1 proteins from 13 species and generated a sequence similarity plot. As well as the canonical UPF1 domains (Fig. 3B, coloured boxes), *Pf*UPF1 has two additional distinct regions: an N-terminal sequence (∼160 residues) that is mostly distinct from the other alveolates (*T. gondii* and *T. thermophila*), and another sequence (∼160 residues) within the UPF2-binding domain that is *Plasmodium*-specific (Fig. 3B, red regions in *P. falciparum* sequence alignment plot). These sequences may be involved in recruiting phylum- or genus-specific proteins to the NMD core complex. However, the key proteins in *P. falciparum* NMD must be able to interact with *Pf*UPF2 in the absence of *Pf*UPF1 and it is possible that because *Pf*UPF2 has derived independence from *Pf*UPF1, these sequence insertions could contribute to *Pf*UPF1 loss-of-function. Given the uncertainty of the role of *Pf*UPF1 in NMD, we next sought to characterise the components of the NMD complex in *P. falciparum*.

**Figure 3.**
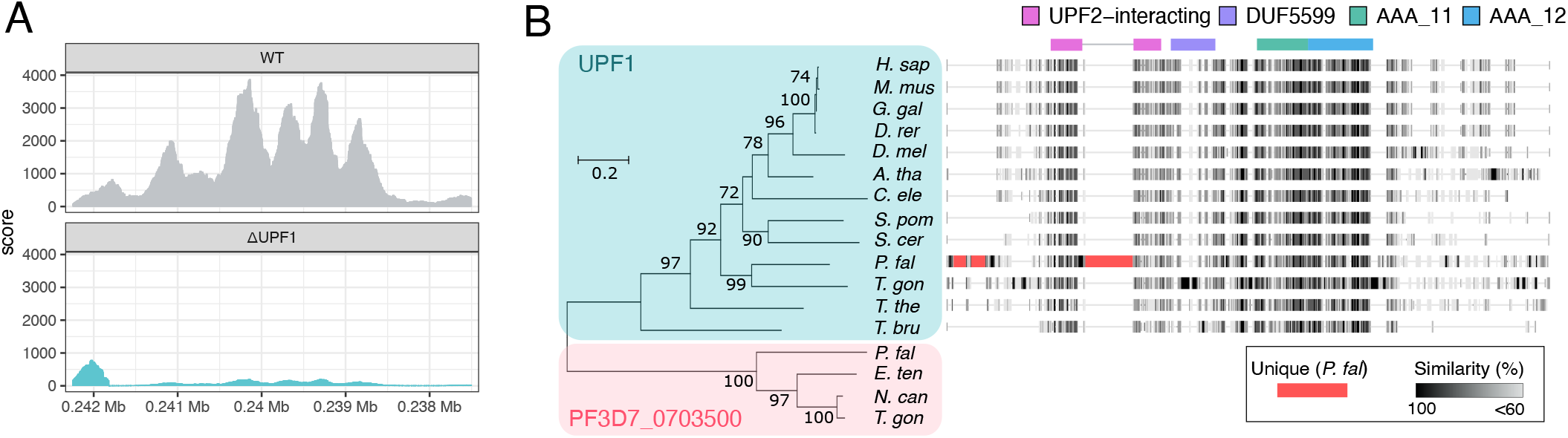
**(A)** Expression of *Pf*UPF1 in WT and Δ*Pf*UPF1 parasites. The coverage plot shows Illumina RNA-seq reads mapped to the PF3D7_1005500 (*Pf*UPF1) locus in WT and Δ*Pf*UPF1 parasites (genomic coordinates Pf3D7_10_v3:237506-242245, negative strand). Each plot represents reads from 3 biological replicates. **(B)** Phylogram created with annotated UPF1 sequences (teal box) and sequences most similar to the PF3D7_0703500 protein identified by BLASTp search (pink box) from the apicomplexans *Eimeria tenella, Neospora caninum* and *T. gondii*. The phylogenetic tree was constructed using the maximum likelihood method and 1000 bootstrap replicates. Bootstrap values <50 are not displayed. Scale bar for the branch length represents the number of substitutions per site. Accession numbers and full alignments are available at https://gitlab.com/e.mchugh/nmd-paper. Conserved UPF1 domains are displayed on top of the sequence alignment. Three longer unique *P. falciparum* sequences are highlighted in red.

### The core NMD proteins co-immunoprecipitate in *P. falciparum*

We used CRISPR-Cas9 as described above with a donor vector encoding a 3X-HA tag to C-terminally epitope tag *Pf*UPF1 and *Pf*UPF2 (Fig. 4A), creating the parasite lines *Pf*UPF1-HA and *Pf*UPF2-HA respectively. Immunofluorescence microscopy of *Pf*UPF1-HA and *Pf*UPF2-HA identified the epitope tag of each protein within the cytoplasm of the parasite, with some fluorescent puncta which may represent processing bodies (P-bodies) (Fig. 4B). P-bodies are conserved ribonucleoprotein granules that contain translationally repressed mRNAs, and mRNA decay factors including proteins involved in NMD (20, 39).

**Figure 4.**
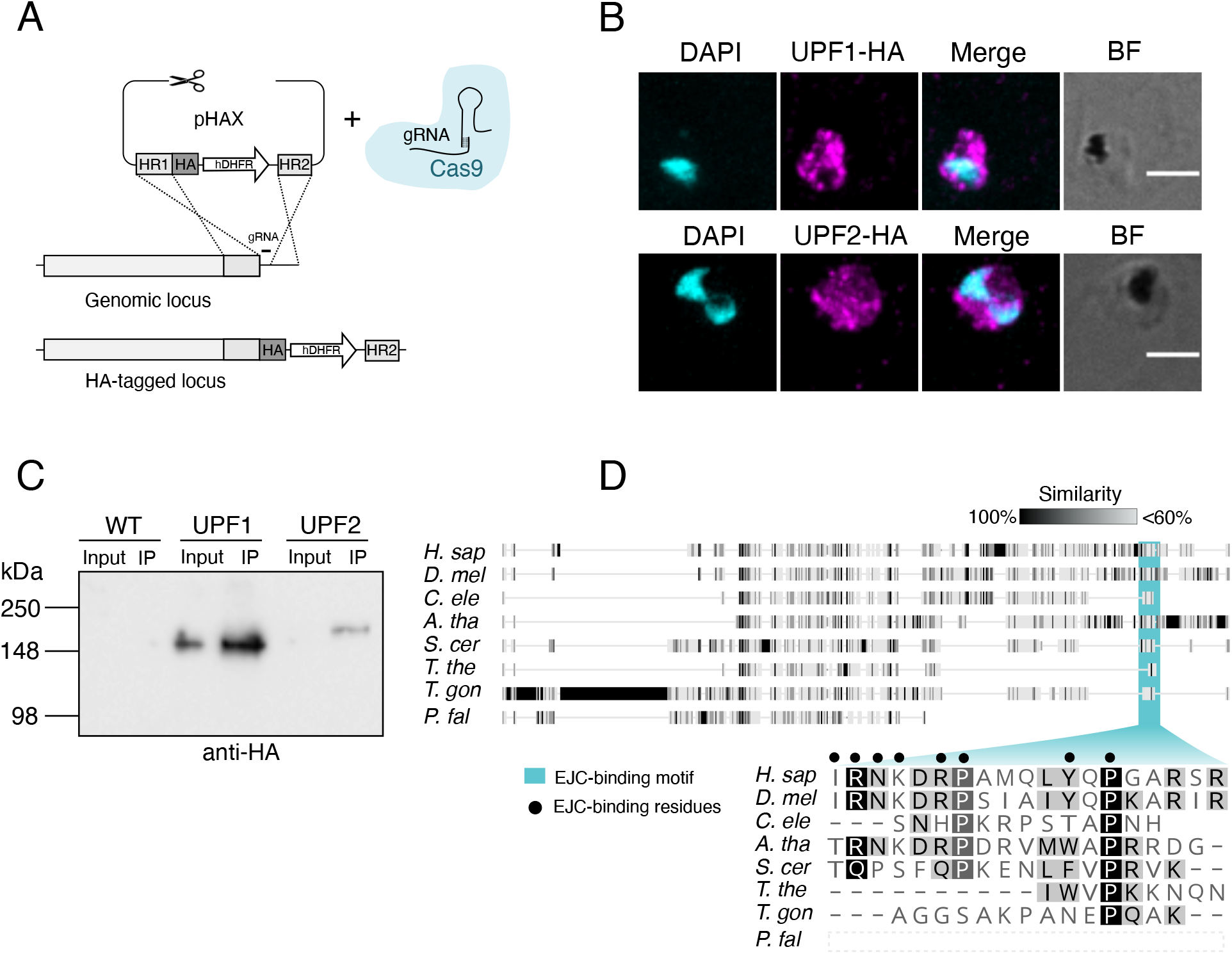
**(A)** Strategy for introducing epitope tags to *Pf*UPF1 and *Pf*UPF2. Homology regions (HRs) targeting the C-terminal of the CDS (excluding the stop codon) and a region in the 3’ UTR were cloned either side of a 3X-HA and hDHFR drug resistance cassette in the plasmid pHAX. **(B)** Immunofluorescence assay with *Pf*UPF1-HA and *Pf*UPF2-HA parasites. Infected RBCs were fixed with paraformaldehyde/glutaraldehyde, permeabilised with Triton X-100 and probed with rat anti-HA (1:300), followed by AlexaFluor anti-rat 568 (1:600). Images are maximum projections of wide-field deconvoluted z-stacks. Nuclei were visualised with DAPI (cyan) and HA signal is presented in magenta. BF = brightfield, scale bars = 3 μm. **(C)** Immunoprecipitation and Western blot of WT, *Pf*UPF1-HA and *Pf*UPF2-HA. For each parasite line, the Input (3% of the Triton X-100-soluble fraction) was loaded beside the immunoprecipitated eluate (100% of the IP). The membrane was probed with anti-HA (1:1000). **(D)** Alignment of UPF3b sequences. The residues in human UPF3b that are important for EJC-binding are denoted with black dots (as determined in (55)). Accession numbers and full alignments are available at https://gitlab.com/e.mchugh/nmd-paper.

We performed co-immunoprecipitation (co-IP) with anti-HA agarose beads of *Pf*UPF1-HA and *Pf*UPF2-HA. Immunoblotting showed that *Pf*UPF1-HA and *Pf*UPF2-HA migrated close to their predicted molecular weights (188 kDa and 212 kDa, respectively) (Fig. 4C). *Pf*UPF1-HA was detected in both the input and IP eluate. Although *Pf*UPF2-HA was successfully enriched and detected in the IP eluate, the concentration in the input was below detection by immunoblotting - likely due to lower expression of *Pf*UPF2. In order to discover proteins that co-IP with *Pf*UPF1-HA and *Pf*UPF2-HA, we performed LC-MS/MS on the IP eluates. Proteins were considered to be co-IPed if they were at least five-fold enriched in the *Pf*UPF1-HA (Table 1) or *Pf*UPF2-HA (Table 2) eluates compared to WT eluate, with a minimum of two significant peptides in both biological replicates. By these criteria, *Pf*UPF2-HA co-IPed with both *Pf*UPF1 and *Pf*UPF3b, indicating that the canonical core NMD proteins interact with one another in *P. falciparum*. *Pf*UPF1-HA did not consistently co-IP with *Pf*UPF2 or *Pf*UPF3b, although peptides from these proteins were detected in one out of two experiments (these proteins of interest are included below the double line in Table 1). As we had identified the putative RNA helicase PF3D7_0703500 by homology to UPF1, we were interested to observe that in the *Pf*UPF2-HA samples, peptides were detected in both replicates, but also in one of the WT controls and hence spectral counts are listed below the double line in Table 2 for interest. In addition to *Pf*UPF1, *Pf*UPF2-HA also co-IPed with another RNA helicase, annotated as DBP1 (PF3D7_0810600). Both *Pf*UPF1-HA and *Pf*UPF2-HA co-IP with a number of ribosomal proteins which is unsurprising as NMD is a translation-dependent process and in other organisms, NMD factors are known to interact directly with some ribosomal components (40).

**Table 1.**
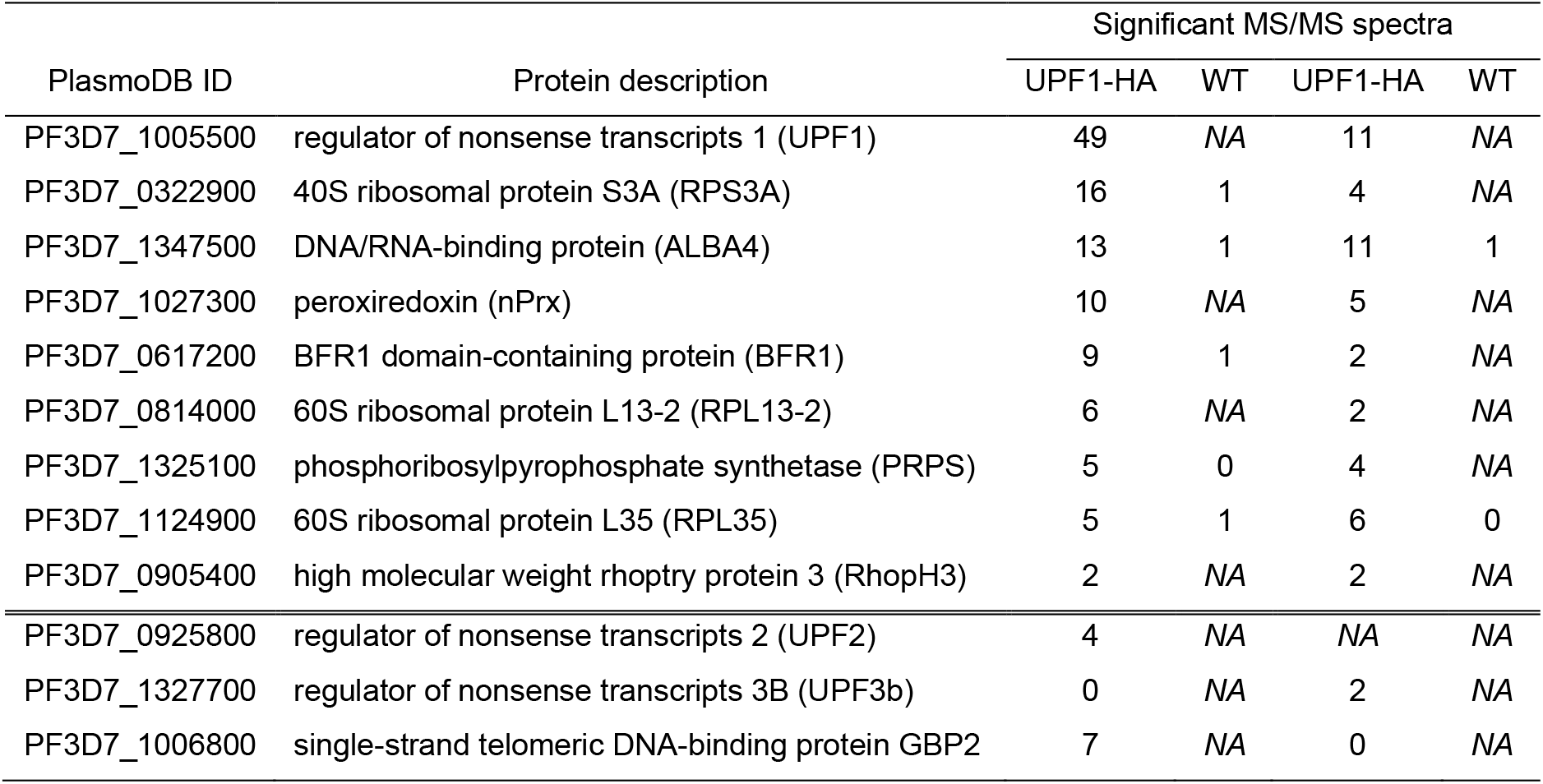
List of proteins co-immunoprecipitated with UPF1-HA

**Table 2.**
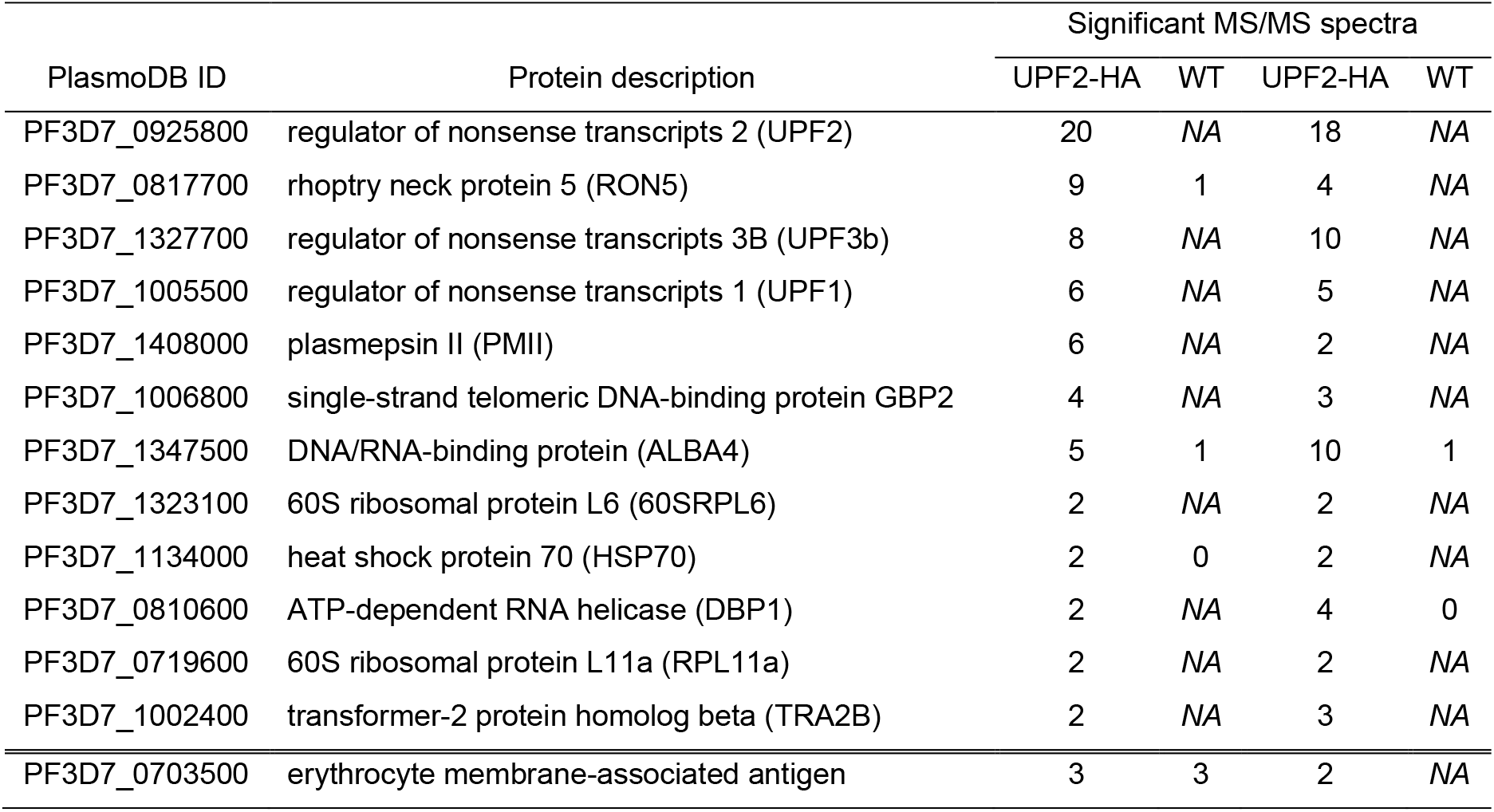
List of proteins co-immunoprecipitated with UPF2-HA

No components of the EJC core complex were identified in any replicate for *Pf*UPF1-HA (Table S5 and Table S6) nor *Pf*UPF2-HA (Table S7 and Table S8). In humans, a conserved motif in UPF3b acts as a bridge between the NMD core and the EJC (41, 42). This EJC-binding motif is absent in *Pf*UPF3b (Fig. 4D, top panel) as this protein is C-terminally truncated compared to human UPF3b (271 vs 483 residues, respectively). The ciliate *T. thermophila*, which performs EJC-independent NMD, also lacks key residues from the human UPF3b EJC-binding motif, (Fig. 4D, bottom panel) (13). It is therefore likely that NMD is also EJC-independent in *P. falciparum*.

We have demonstrated that NMD occurs (albeit divergent from classical NMD) and that the core NMD interacting complex is conserved in *P. falciparum*. However, it is unclear how the NMD core complex couples PTC-containing transcripts to RNA decay machinery, such as nucleases or the exosome. These steps in NMD are not conserved and are poorly understood outside of metazoans. We were therefore interested in identifying RNA-binding proteins that interact with the NMD core complex that may be involved in this process. A network of all protein interactions identified by co-IP is presented in Fig. 5A. Of particular interest is an apicomplexan-specific RNA-binding protein *Pf*ALBA4 (PF3D7_1347500), that co-IPed with both *Pf*UPF1-HA and *Pf*UPF2-HA. In asexual *Plasmodium yoelii*, disruption of *Py*ALBA4 led to an increase in mRNA abundance, which may imply a role for ALBA4 in mRNA decay (43). We found that *Pf*UPF2-HA also co-IPed with another putative RNA-binding protein, *Pf*GBP2 (G-strand binding protein 2; PF3D7_1006800). This protein was also identified in one out of two of the *Pf*UPF1-HA co-IP experiments (Table 1, below double line). A study in *P. yoelii* showed that *Py*ALBA4 co-IPed with *Py*GBP2, which was one of the most abundant interacting proteins (Fig. 5A, dotted line) (43). This suggests that interactions between UPF2, ALBA4 and GBP2 are conserved in *Plasmodium* (43), however, the function of *Pf*GBP2 has not been studied. In yeast, Gbp2 is an SR (serine-arginine) protein that marks unspliced transcripts and targets them for elimination via the nuclear exosome (44). However, it seems unlikely that *Pf*GBP2 functions in exactly the same way, as the RS (serine-arginine repeat) domain is not conserved, and unlike in yeast, the *P. falciparum* RNA exosome is mainly cytoplasmic (45). Yeast Gbp2 also recruits nuclear export protein Mex67 to correctly spliced transcripts and shuttles to the cytoplasm, although in the more closely related apicomplexan *T. gondii, Tg*GBP2 (TgRRM_2620) is not required for nuclear export of mRNA and *P. falciparum* lacks the nuclear export protein (Mex67) that interacts with GBP2 (46). Another study in yeast found that Gbp2 associated with Upf1 (12). We speculate that when exported to the cytoplasm, *Pf*GBP2 is a marker of unspliced, intron-containing transcripts that serves as part of an NMD-stimulating environment on target mRNAs. Other putative splicing proteins that may remain on exported mRNAs, such as TRA2B (PF3D7_1002400; co-IPed with *Pf*UPF2-HA) could also be involved in triggering NMD. We also identified *Pf*BFR1 (a homologue of the yeast protein brefeldin A resistance factor 1; Brf1p) in the co - IP eluate of *Pf*UPF1. In yeast, Bfr1p is an RNA-binding protein that targets mRNAs to cytoplasmic sites of mRNA decay called P-bodies (47). Characterisation of P-body-associated proteins and other RNA-binding proteins that interact with the NMD core in *P. falciparum* will be important for elucidating the process of NMD in this parasite. Based on our co-IP and RNA-seq findings, we suggest a model for NMD in *P. falciparum* which centres on *Pf*UPF2 recognising markers of unspliced introns after ribosome stalling at a PTC (Fig. 5B). One shortcoming of this model is that we do not know if NMD occurs for PTC-containing transcripts arising from processes other than intron retention, for example from transcriptional error, and our model does not explain how these mRNAs might be identified and degraded.

**Figure 5.**
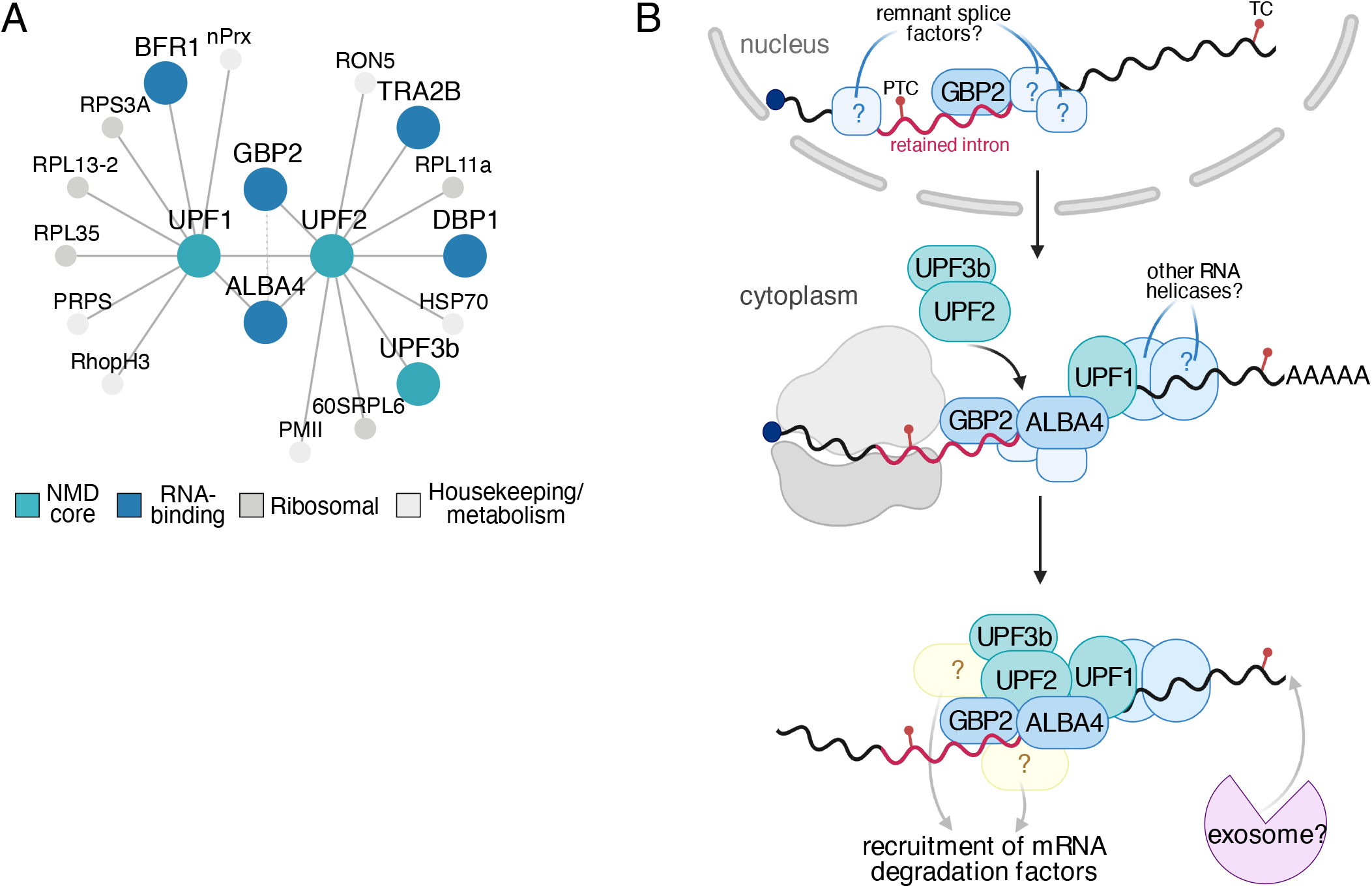
**(A)** Map of *Pf*UPF1-HA and *Pf*UPF2-HA protein-protein interactions detected by co-IP and LC-MS/MS. Dashed line represents interaction detected in a co-immunoprecipitation experiment with *Py*ALBA4-GFP in *P. yoelii* in (56). Full protein names are listed in Tables 1 and 2. **(B)** Model of NMD in *P. falciparum* based on protein-protein interactions detected in this study. During translation, the ribosome stalls on a PTC introduced by intron retention. In this model of NMD, the splicing factor *Pf*GBP2 remains on an unspliced intron after mRNA export to the cytoplasm. *Pf*GBP2, and the RNA-binding protein *Pf*ALBA4, may aid in PTC-containing transcript recognition by the NMD factor *Pf*UPF2. *Pf*UPF2 interacts with the other NMD core complex members *Pf*UPF1 and *Pf*UPF3b, leading to the initiation of NMD and degradation of the transcript through an as yet unidentified mechanism.

This study describes a highly divergent, UPF1-independent NMD pathway. Our findings corroborate other work (34) that suggests no special regulatory role for the majority of observed intron retention. Further work is needed to identify specific NMD substrates and NMD accessory proteins in order to understand how NMD has evolved in the malaria parasite.

## MATERIALS AND METHODS

### Molecular biology and transfection of *Plasmodium falciparum*

Parasite lines with endogenously HA-tagged putative NMD components, were generated by CRISPR using the episomal expression plasmid pHAX as a donor template. To generate pHAX, DNA sequence encoding a 3xHA tag was amplified from pGLMS-HA (48) using 3HA-FOR (GCG**ACGCGTG**CTTACCCGTACGACGTC) and 3HA-REV (GCG**TTAATTAA** TTAAGCAGCGGCATAATCTGG) primers (MluI and PacI in bold). pGLUX1-PfCentrin2-mCherry (unpublished) was digested with MluI and PacI to release the mCherry and the PCR product was directionally cloned into the to generate pHAX-*Pf*Centrin2. The sequence map for pHAX-*Pf*Centrin2 is supplied in the Supplementary data files.

For CRISPR-Cas9 editing, guide RNA (gRN A) binding sites with a protospacer adjacent motif (PAM) were selected for PF3D7_ 1005500 (*Pf*UPF1) and PF3D7_0925800 (*Pf*UPF2) using CHOPCHOP (https://chopchop.cbu.uib.no/) and were synthesised as crRNA (Integrated DNA Technologies) (49). Homology regions (HRs) were PCR amplified from NF54 genomic DNA with Phusion polymerase (NEB) using primers listed in Table S9. For gene disruption, two homology regions (HRs) 450-630 bp in length were inserted either side of the human dihydrofolate reductase (hDHFR) drug selection cassette in pGLMS-HA (48) at the BglII and XhoI sites (HR1), and EcoRI and KasI (HR2) using the In-Fusion®HD Cloning Kit (Takara Bio). For C-terminal HA-tagging, HRs were inserted at the XhoI and MluI sites (HR1), and NcoI and KasI sites (HR2) in pHAX-*Pf*Centrin2.

Homology regions were no more than 30 bp from the predicted site of the Cas9-induced DNA double-strand break. Plasmids were confirmed by Sanger sequencing (AGRF Melbourne). Final plasmids (100 μg) were linearised with BglII and BglI then precipitated and resuspended in cytomix.

Linear templates were endogenously integrated using commercial Cas9 nuclease and synthesised gRNA. To our knowledge, this is a novel technique in transfection of *P. falciparum* and has some benefits over plasmid expression systems, as it does not require molecular cloning and plasmid preparation for Cas9 and guide components. Prior to transfection, gene-specific crRNA (100 μM) and tracrRNA (100 μM) were annealed to form gRNA (Integrated DNA Technologies) by mixing 1:1, heating to 95°C for 5 min and allowing to cool to room temperature. The gRNA (3 μL) was then complexed with 2 μL *Streptococcus pyogenes* Cas9 nuclease (Integrated DNA Technologies) at room temperature for 20 min. The resulting ribonucleoprotein (5 μL) was added to the linear DNA template and electroporated into ring stage NF54 infected RBCs as previously described (50). Parasites resistant to 5 nM WR99210 (Jacobus Pharmaceuticals) were observed by Giemsa smear 13-21 days following transfection. Genomic DNA was extracted from wildtype NF54, Δ*Pf*UPF1 and Δ*Pf*UPF2 parasite cultures using QuickExtract (Lucigen). The identity of transfectant parasites were verified by PCR specific for the expected parental and modified loci. PCR was performed with GoTaq (Promega) and with primers listed in Table S9.

### Illumina RNA sequencing

Parasite cultures were synchronised with 5% (w/v) D-sorbitol in H_2_O in the cycle before harvest (51). Trophozoite-stage parasites (∼10^10^ parasites) were isolated with 0.03% (w/v) saponin in PBS, washed with PBS and then resuspended in 1 mL TRI Reagent (Sigma Aldrich). Chloroform (200 μL) was added to the parasite/TRI Reagent sample and was mixed by vortex for 15 s, incubated for 3 min at room temperature, then centrifuged at 12,000 *g* for 30 min at 4°C. The aqueous supernatant was removed and concentrated using a RNeasy MinElute kit (Qiagen). Samples were then treated with DNase I (Qiagen) and concentrated again using RNeasy MinElute columns (Qiagen). Library preparation and sequencing was performed by Victorian Clinical Genetics Services using TruSeq stranded mRNA kit (Illumina) and sequenced on a NovaSeq 6000 to a depth of 30M reads per sample, reads were 150 base pairs, paired-end reads with three biological replicates for each sample.

### Bioinformatics analyses

RNA-seq read quality was checked using FastQC (0.11.8) before mapping of reads to the *P. falciparum* 3D7 genome (release 45) using STAR (2.7.3) (52, 53). A summary of our RNA-seq read mapping is presented in Table S10. Differential expression of genes was tested with limma-voom using the *treat* method to determine adjusted p-values (requiring a log-fold change of at least 1). Genes with adjusted p-values < 0.05 were considered differentially expressed. Read counts used for calculating intron retention, alternative splicing analysis, including differential alternative splicing were performed using ASpli (1.10.0). Gene expression FPKM values were determined using RSeqQC (FPKM_count.py). The introduction of premature termination codons following intron retention was determined using a purpose-written R script available at gitlab.com/e.mchugh/nmd-paper. All plots were generated in R using ggplot2 (3.3.3). Protein schema were generated using the R package drawProteins (1.9.1) (54). Coverage plots were created using the R package superintronic (0.99.4) (55). Analyses were performed using the Spartan High Performance Computing system (University of Melbourne) or on personal computers. Commands and scripts used are available at gitlab.com/e.mchugh/nmd-paper. RNA sequencing data files are available on the NCBI Sequence Read Archive with the BioProject identifier PRJNA699307.

### Phylogenetic tree construction

UPF1 and UPF3b protein sequences were obtained through OrthoMCL (https://orthomcl.org/). PF3D7_0703500-related sequences were obtained by BLASTp search (https://blast.ncbi.nlm.nih.gov/) using the PF3D7_0703500 protein sequence as input and restricting results to exclude the *Plasmodium* taxon. Multiple sequence alignment was performed using the MAFFT-linsi method (v7.475) (56) and a sequence similarity plot was generated using Geneious Prime (2019.1.3) (57). Alignments were trimmed using trimAl (v1.3) (58) before construction of a maximum-likelihood phylogenetic tree with 1,000 bootstrap replicates and phylogram plotting using MEGAX (10.1.0) (59).

### Growth analysis

Cultures were initiated at 1% parasitaemia, 1% haematocrit in triplicate. Parasitaemia was measured by flow cytometer (BD FACSCanto) every 48 h by staining with SYTO-61 (Invitrogen). Identical subculturing was performed on wildtype and knockout parasites to prevent overgrowth. Flow cytometry data were analysed using FlowJo™ (10.6.0) software.

### Immunofluorescence microscopy

Infected RBCs were harvested and resuspended in PBS at 5% haematocrit. Coverslips were coated with lectin (from *Phaseolus vulgaris* ; PHA-E, Sigma product L8629), washed, and infected RBCs were applied. Adhered cells were washed 3X with PBS until a monolayer remained on the coverslip. The cells were then fixed for 20 min in 4% paraformaldehyde/0.008% glutaraldehyde in PBS, followed by permeabilisation with 0.1% Triton X-100 in PBS for 10 min. Rat anti-HA antibody (1:300) in 3% (w/v) BSA in PBS was applied for 1.5 h, washed 3X with PBS, followed by incubation with anti-rat AlexaFluor 568 (1:600) for 1 h. The nuclear stain DAPI (4′,6-diamidino-2-phenylindole) was added prior to mounting and sealing of the slide with nailpolish. Slides were imaged on a GE DeltaVision Elite Widefield Deconvolution Microscope. Deconvolved images were processed using FIJI software (60). Images stacks are presented as maximum projections and have been adjusted by cropping, adding false colour and changes to brightness/contrast.

### Co-immunoprecipitation, immunoblotting and mass spectrometry

Infected RBCs were synchronised with 5% (w/v) D-sorbitol in H_2_O and parasites were harvested as trophozoites 72 h later. Parasites were isolated from RBCs by lysis with 0.03% (w/v) saponin in PBS on ice. Parasite pellets were then solubilised in immunoprecipitation buffer (IP buffer) containing 1% Triton X-100, 50 mM Tris-HCl, 150 mM NaCl, 2 mM EDTA and cOmplete ™ EDTA-free Protease Inhibitor Cocktail (Roche product 11836170001) for 30 min on ice. Insoluble material was separated by centrifugating twice at 13,000 *g* for 10 min. The Triton X-100-soluble fraction (input) was incubated with anti-HA agarose beads (Roche product ROAHAHA) overnight at 4°C and was washed five times with IP buffer. For immunoblotting, proteins were eluted at 95 °C for 10 min with Laemmli buffer containing β-mercaptoethanol. Input and eluate samples were separated on 4-15% Tris-glycine polyacrylamide gels at 200V for 35 min. For mass spectrometry, beads wer e washed a further two times in 1 mM Tris-HCl (pH 7.4) before elution with 0.1 % (v/v) formic acid and trifluoroethanol at 50 °C for 5 min. The eluate was neutralised with triethylammonium bicarbonate and reduced with TCEP (5 mM), then digested with trypsin for 16 h at 37°C. Samples were then analysed by LC-MS/MS with a Q Exactive Plus mass spectrometer. Mass spectra were searched using MASCOT against a custom protein database comprised of the 3D7 *P. falciparum* annotated proteome (version 43) and *Homo sapiens* reference proteome (Uniprot proteome ID: UP000005640). MASCOT searches were performed with the following parameters: MS tolerance = 10 ppm, MS/MS tolerance = 0.2 Da, cleavage enzyme = trypsin, missed cleavages allowed = 3, peptide isotope error = 0, variable modifications = oxidation (M), with a MASCOT decoy search performed concurrently. Proteins were considered to be enriched in *Pf*UPF1-HA or *Pf*UPF2-HA IP eluate compared to a wildtype control if there were at least two significant peptides detected in two biological replicates and a) five-fold the number of significant peptides detected or b) no peptides were detected in the control. False discovery rates for each experiment are presented in Table S11. The mass spectrometry proteomics data have been deposited to the ProteomeXchange Consortium via the PRIDE partner repository with the dataset identifier PXD023910.

## AVAILABILITY

Computer scripts used for analysis of PTCs, statistical analysis and generating figures are available at https://gitlab.com/e.mchugh/nmd-paper.

## Supporting information

pHAX-PfCentrin2

## ACCESSION NUMBERS

Illumina RNA-seq data have been deposited in the NCBI Sequence Read Archive BioProject under accession number PRJNA699307. Co-immunoprecipitation LC-MS/MS data are available in the PRIDE repository under accession number PXD023910.

## SUPPLEMENTARY DATA

Supplementary data are included separately.

## ACKNOWLEDGEMENTS

Red blood cells were provided Australian Red Cross Lifeblood. Thank you to Dr. Lee Yeoh and Dr. Laurent Duret for helpful discussions on bioinformatics approaches. The authors also thank Dr. Simon Cobbold for his advice on CRISPR-Cas9.

---

**Figure S1.**
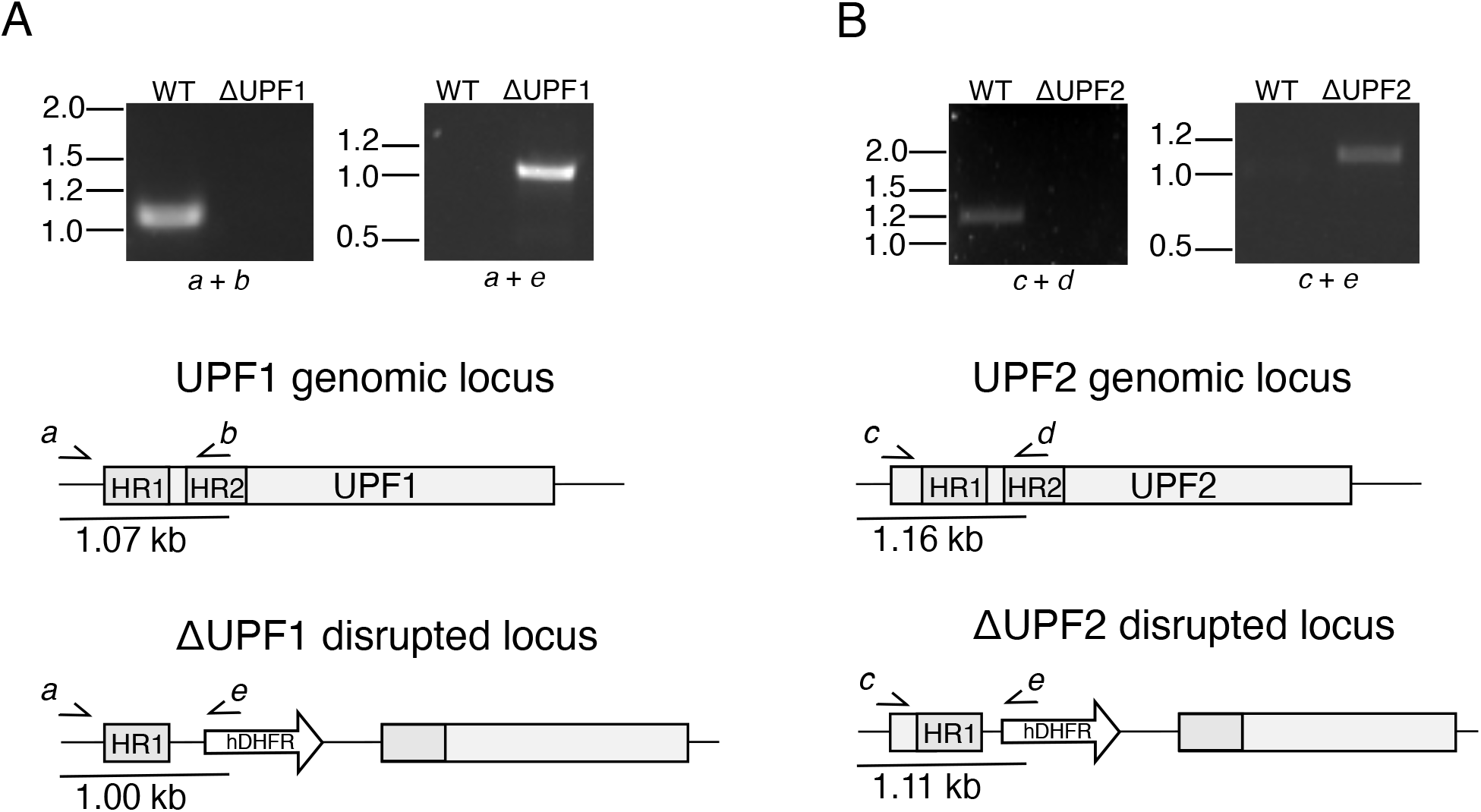
PCR confirmation of gene disruption in Δ*Pf*UPF1 and Δ*Pf*UPF2 parasites. **(A)** A PCR product (1.07 kb, primers *a* + *b*) is amplified from the *Pf*UPF1 locus with WT parasite DNA but not Δ *Pf*UPF1 parasite DNA. An amplicon (1.00 kb, primers *a* + *e*) from the disrupted Δ*Pf*UPF1 locus, resulting from integration of the hDHFR cassette, is amplified only from Δ*Pf*UPF1 parasite DNA. **(B)** An amplicon (1.16 kb, primers *c* + *d*) resulting from the *Pf*UPF2 genomic locus is only amplified from WT parasite DNA. Note that primer ‘*c*’ anneals upstream of the 5’ homology region. A product (1.11 kb, primers *c* + *e*) amplified from only Δ *Pf*UPF2 parasite DNA confirms disruption of the CDS with the hDHFR cassette. Primer sequences are listed in Table S9.

**Figure S2.**
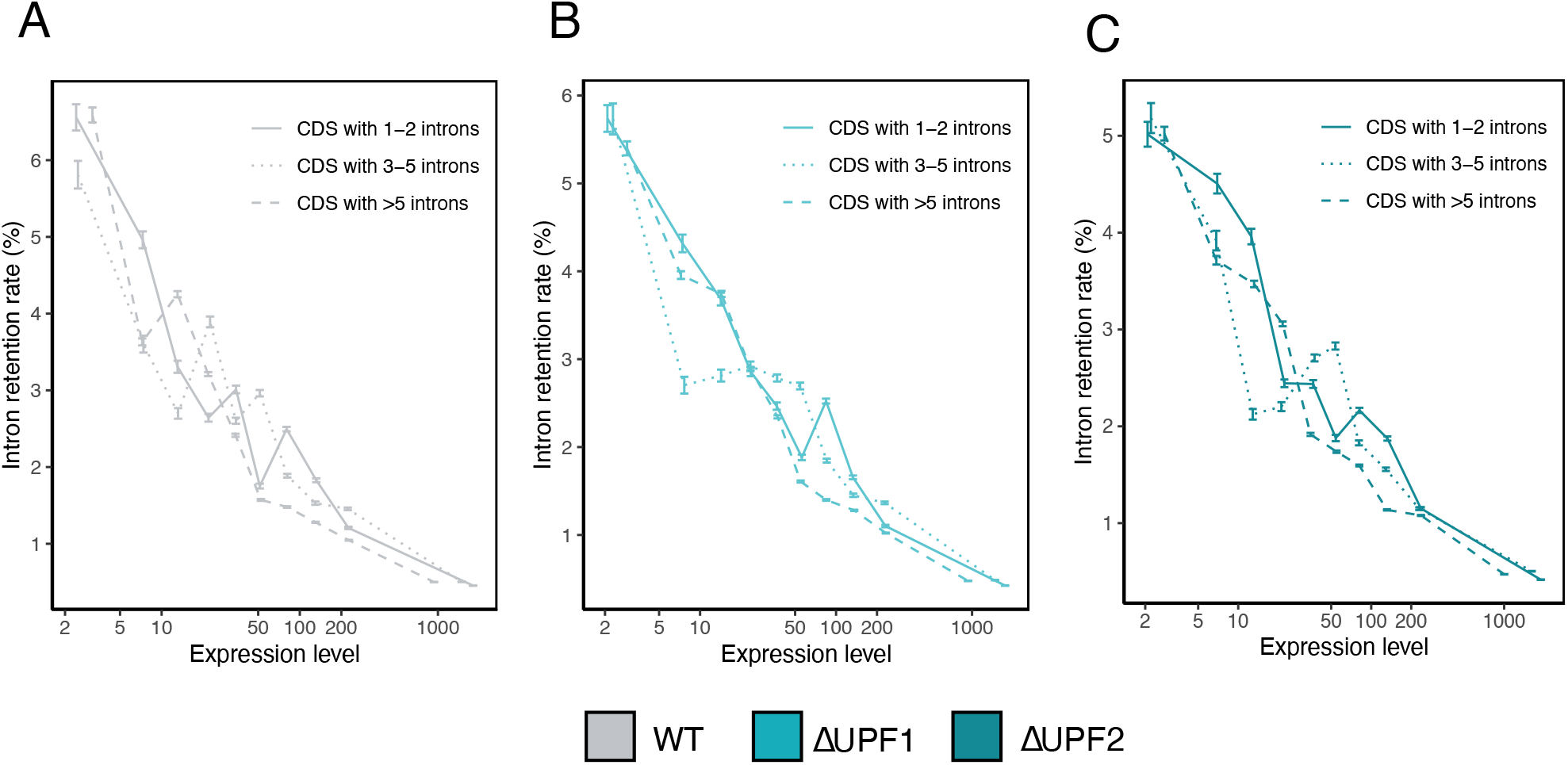
Introns (n = 7523) were grouped into 10 equal-sized bins based on expression level (FPKM) and for **(A)** WT and **(B)** Δ *Pf*UPF1 and **(C)** Δ*Pf*UPF2 the IR rate was computed globally within each bin as a proportion. CDS 1-2 = 1969 introns, CDS with 3-5 = 2222 introns, CDS with >5 introns = 3332. Error bars represent 95% confidence interval.

**Figure S3.**
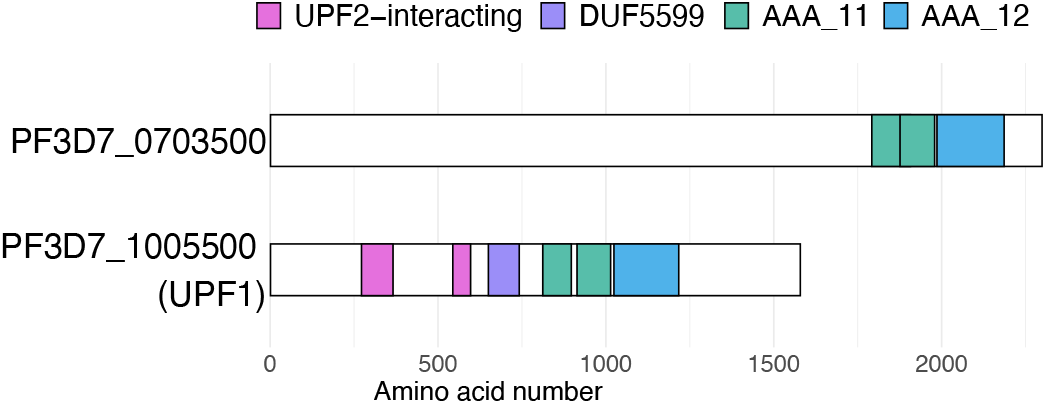
Protein schema showing Pfam domains in PF3D7_0703500 and *Pf*UPF1. AAA_11 and AAA_12 are AAA ATPase domains associated with ATP-dependent helicase activity. DUF6699 (DUF = domain of unknown function) is a domain also present in the *Homo sapiens* UPF1. The N-terminal UPF2-interacting domain contains zinc binding motifs which interact with UPF2.

**Table S9.**
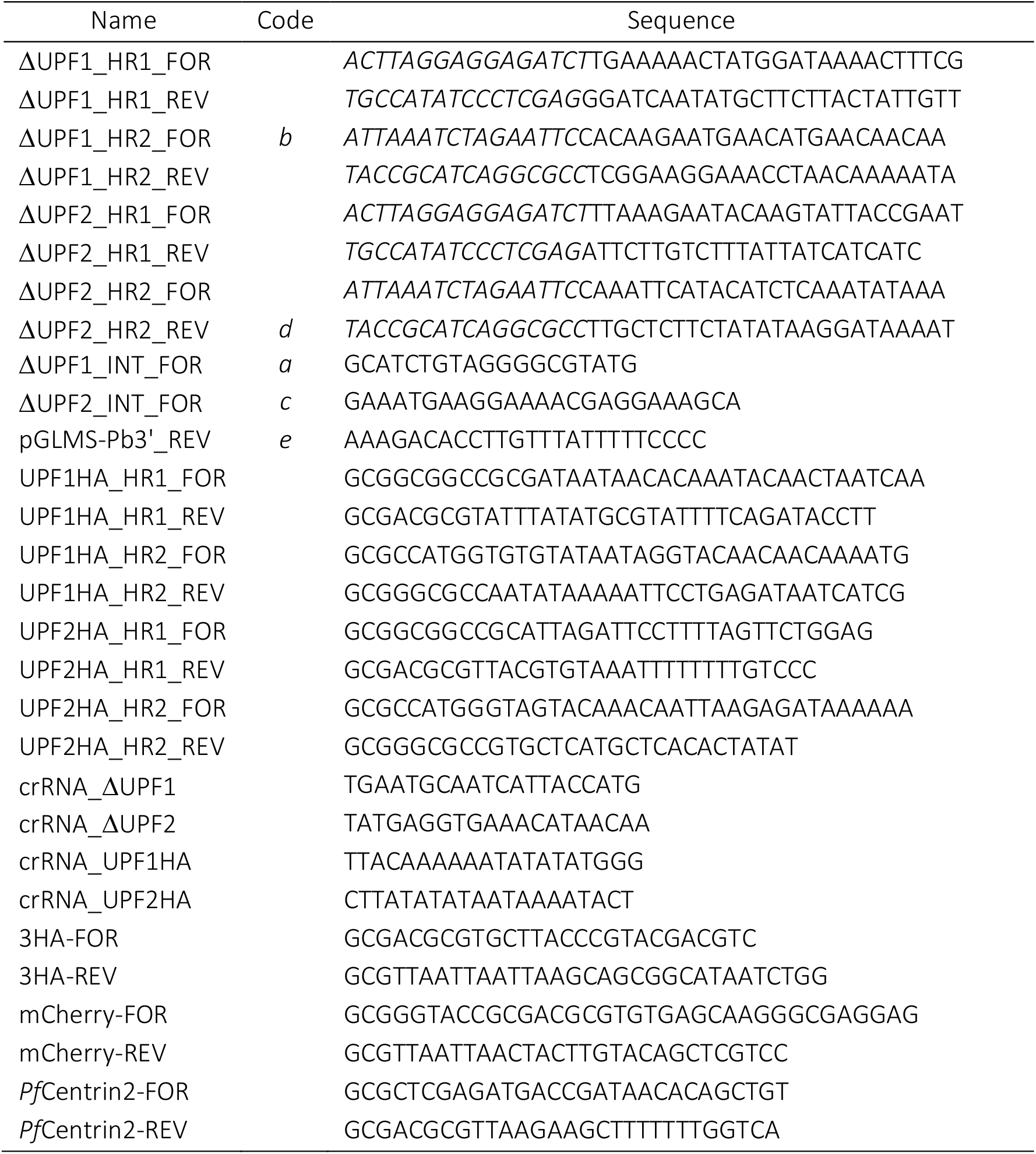
Oligonucleotides used in this study. Italics indicates sequences homologous to vector.

**Table S10.** Illumina STAR mapping information.

| Sample | Replicate | Input reads | Uniquely mapped (%) | Average input<br>read length | FASTQ accession (SRA) |
| --- | --- | --- | --- | --- | --- |
| NF54 | 1 | 40823544 | 85.97 | 291 | SRR13622007 |
| NF54 | 2 | 46313559 | 89.61 | 291 | SRR13622006 |
| NF54 | 3 | 32861972 | 92.10 | 291 | SRR13622014 |
| ΔUPF1 | 1 | 24369887 | 88.50 | 293 | SRR13622013 |
| ΔUPF1 | 2 | 48948306 | 93.53 | 290 | SRR13622012 |
| ΔUPF1 | 3 | 33797287 | 90.46 | 291 | SRR13622011 |
| ΔUPF2 | 1 | 30679940 | 89.08 | 295 | SRR13622010 |
| ΔUPF2 | 2 | 29202646 | 86.51 | 293 | SRR13622009 |
| ΔUPF2 | 3 | 30632528 | 88.49 | 293 | SRR13622008 |

**Table S11.** False discovery rates for mass spectrometry MASCOT searches for peptide matches above the identity threshold.

| Experiment | False discovery rate |
| --- | --- |
| NF54_13-11-20 | 1.70% |
| UPF1_13-11-20 | 0.96% |
| NF54_17-11-20 | 1.75% |
| UPF2_17-11-20 | 2.22% |
| NF54_30-11-20 | 2.29% |
| UPF1_30-11-20 | 1.63% |
| UPF2_30-11-20 | 2.91% |

## REFERENCES

1. Thermann R, Neu-Yilik G, Deters A, Frede U, Wehr K, Hagemeier C, et al. Binary specification of nonsense codons by splicing and cytoplasmic translation. The EMBO journal. 1998;17(12):3484–94.

2. Lykke-Andersen J, Shu MD, Steitz JA. Communication of the position of exo n-exon junctions to the mRNA surveillance machinery by the protein RNPS1. Science. 2001;293(5536):1836–9.

3. Amrani N, Ganesan R, Kervestin S, Mangus DA, Ghosh S, Jacobson A. A faux 3’ -UTR promotes aberrant termination and triggers nonsense-mediated mRNA decay. Nature. 2004;432(7013):112–8.

4. Hamid FM, Makeyev EV. Emerging functions of alternative splicing coupled with nonsense - mediated decay. Biochemical Society transactions. 2014;42(4):1168–73.

5. Nickless A, Bailis JM, You Z. Control of gene expression through the nonsense-mediated RNA decay pathway. Cell Biosci. 2017;7:26.

6. Lareau LF, Brenner SE. Regulation of splicing factors by alternative splicing and NMD is conserved between kingdoms yet evolutionarily flexible. Molecular biology and evolution. 2015;32(4):1072–9.

7. Wong JJ, Ritchie W, Ebner OA, Selbach M, Wong JW, Huang Y, et al. Orchestrated intron retention regulates normal granulocyte differentiation. Cell. 2013;154(3):583–95.

8. Celik A, Baker R, He F, Jacobson A. High -resolution profiling of NMD targets in yeast reveals translational fidelity as a basis for substrate selection. RNA. 2017;23(5):735–48.

9. Raxwal VK, Simpson CG, Gloggnitzer J, Entinze JC, Guo W, Zhang R, et al. Nonsense - Mediated RNA Decay Factor UPF1 Is Critical for Posttranscriptional and Translational Gene Regulation in Arabidopsis. Plant Cell. 2020;32(9):2725–41.

10. Lee VV, Judd LM, Jex AR, Holt KE, Tonkin CJ, Ralph SA. Direct Nanopore Sequencing of mRNA Reveals Landscape of Transcript Isoforms in Apicomplexan Parasites. mSystems. 2021;6(2).

11. Chen YH, Su LH, Sun CH. Incomplete nonsense -mediated mRNA decay in Giardia lamblia. International journal for parasitology. 2008;38(11):1305–17.

12. Dehecq M, Decourty L, Namane A, Proux C, Kanaan J, Le Hir H, et al. Nonsense -mediated mRNA decay involves two distinct Upf1 -bound complexes. The EMBO journal. 2018;37(21).

13. Tian M, Yang W, Zhang J, Dang H, Lu X, Fu C, et al. Nonsense-mediated mRNA decay in Tetrahymena is EJC independent and requires a protozoa-specific nuclease. Nucleic acids research. 2017;45(11):6848–63.

14. Okada-Katsuhata Y, Yamashita A, Kutsuzawa K, Izumi N, Hirahara F, Ohno S. N - nd Cterminal Upf1 phosphorylations create binding platforms for SMG-6 and SMG-5:SMG-7 during NMD. Nucleic acids research. 2012;40(3):1251–66.

15. Causier B, Li Z, De Smet R, Lloyd JPB, Van de Peer Y, Davies B. Conservation of Nonsense - Mediated mRNA Decay Complex Components Throughout Eukaryotic Evolution. Scientific reports. 2017;7(1):16692.

16. Treeck M, Sanders JL, Elias JE, Boothroyd JC. The phosphoproteomes of Plasmodium falciparum and Toxoplasma gondii reveal unusual adaptations within and beyond the parasites’ boundaries. Cell Host Microbe. 2011;10(4):410–9.

17. Mair GR, Lasonder E, Garver LS, Franke-Fayard BM, Carret CK, Wiegant JC, et al. Universal features of post-transcriptional gene regulation are critical for Plasmodium zygote development. PLoS Pathog. 2010;6(2):e1000767.

18. Yeoh LM, Goodman CD, Mollard V, McHugh E, Lee VV, Sturm A, et al. Alternative splicing is required for stage differentiation in malaria parasites. Genome Biol. 2019;20(1):151.

19. Lunghi M, Spano F, Magini A, Emiliani C, Carruthers VB, Di Cristina M. Alternative splicing mechanisms orchestrating post-transcriptional gene expression: intron retention and the intron-rich genome of apicomplexan parasites. Curr Genet. 2016;62(1):31–8.

20. Reddy BP, Shrestha S, Hart KJ, Liang X, Kemirembe K, Cui L, et al. A bioinformatic survey of RNA-binding proteins in Plasmodium. BMC Genomics. 2015;16:890.

21. Bryant JM, Baumgarten S, Glover L, Hutchinson S, Rachidi N. CRISPR in Parasitology: Not Exactly Cut and Dried! Trends in parasitology. 2019;35(6):409–22.

22. Volz JC, Yap A, Sisquella X, Thompson JK, L im NT, Whitehead LW, et al. Essential Role of the PfRh5/PfRipr/CyRPA Complex during Plasmodium falciparum Invasion of Erythrocytes. Cell Host Microbe. 2016;20(1):60–71.

23. Spillman NJ, Beck JR, Ganesan SM, Niles JC, Goldberg DE. The chaperonin TRiC forms an oligomeric complex in the malaria parasite cytosol. Cell Microbiol. 2017;19(6).

24. Ghorbal M, Gorman M, Macpherson CR, Martins RM, Scherf A, Lopez-Rubio JJ. Genome editing in the human malaria parasite Plasmodium falciparum using the CRISPR-Cas9 system. Nat Biotechnol. 2014;32(8):819–21.

25. Cho SW, Lee J, Carroll D, Kim JS, Lee J. Heritable gene knockout in Caenorhabditis elegans by direct injection of Cas9-sgRNA ribonucleoproteins. Genetics. 2013;195(3):1177–80.

26. Woo JW, Kim J, Kwon SI, Corvalan C, Cho SW, Kim H, et al. DNA-free genome editing in plants with preassembled CRISPR-Cas9 ribonucleoproteins. Nat Biotechnol. 2015;33(11):1162–4.

27. Sung YH, Kim JM, Kim HT, Lee J, Jeon J, Jin Y, et al. Highly efficient gene knockout in mice and zebrafish with RNA-guided endonucleases. Genome research. 2014;24(1):125–31.

28. Kim S, Kim D, Cho SW, Kim J, Kim JS. Highly efficient RNA-guided genome editing in human cells via delivery of purified Cas9 ribonucleoproteins. Genome research. 2014;24(6):1012–9.

29. Schumann K, Lin S, Boyer E, Simeonov DR, Subramaniam M, Gate RE, et al. Generation of knock-in primary human T cells using Cas9 ribonucleoproteins. Proc Natl Acad Sci U S A. 2015;112(33):10437–42.

30. Zhang M, Wang C, Otto TD, Oberstaller J, Liao X, Adapa SR, et al. Uncovering the essential genes of the human malaria parasite Plasmodium falciparum by saturation mutagenesis. Science. 2018;360(6388).

31. Lareau LF, Inada M, Green RE, Wengrod JC, Brenner SE. Unproductive splicing of SR genes associated with highly conserved and ultraconserved DNA elements. Nature. 2007;446(7138):926–9.

32. Guizetti J, Scherf A. Silence, activate, poise and switch! Mechanisms of antigenic variation in Plasmodium falciparum. Cell Microbiol. 2013;15(5):718 –26.

33. Braunschweig U, Barbosa-Morais NL, Pan Q, Nachman EN, Alipanahi B, Gonatopoulos -x Pournatzis T, et al. Widespread intron retention in mammals functionally tunes transcriptomes. Genome research. 2014;24(11):1774–86.

34. Saudemont B, Popa A, Parmley JL, Rocher V, Blugeon C, Necsulea A, et al. The fitness cost of mis-splicing is the main determinant of alternative splicing patterns. Genome Biol. 2017;18(1):208.

35. Delhi P, Queiroz R, Inchaustegui D, Carrington M, Clayton C. Is there a classical nonsense - mediated decay pathway in trypanosomes? PloS one. 2011;6(9):e25112.

36. Fourati Z, Roy B, Millan C, Coureux PD, Kervestin S, van Tilbeurgh H, et al. A highly conserved region essential for NMD in the Upf2 N -terminal domain. Journal of molecular biology. 2014;426(22):3689–702.

37. Kun J, Hesselbach J, Schreiber M, Scherf A, Gysin J, Mattei D, et al. Cloning and expression of genomic DNA sequences coding for putative erythrocyte membrane -associated antigens of Plasmodium falciparum. Res Immunol. 1991;142(3):199–210.

38. Bunnik EM, Batugedara G, Saraf A, Prudhomme J, Florens L, Le Roch KG. The mRNA -bound proteome of the human malaria parasite Plasmodium falciparum. Genome Biol. 2016;17(1):147.

39. Park Y, Park J, Hwang HJ, Kim B, Jeong K, Chang J, et al. Nonsense -mediated mRNA decay factor UPF1 promotes aggresome formation. Nat Commun. 2020;11(1):3106.

40. Gupta P, Li YR. Upf proteins: highly conserved factors involved in nonsense mRNA mediated decay. Mol Biol Rep. 2018;45(1):39 –55.

41. Kunz JB, Neu-Yilik G, Hentze MW, Kulozik AE, Gehring NH. Functions of hUpf3a and hUpf3b in nonsense-mediated mRNA decay and translation. RNA. 2006;12(6):1015 –22.

42. Buchwald G, Ebert J, Basquin C, Sauliere J, Jayachandran U, Bono F, et al. Insights into the recruitment of the NMD machinery from the crystal structure of a core EJC-UPF3b complex. Proc Natl Acad Sci U S A. 2010;107(22):10050–5.

43. Munoz EE, Hart KJ, Walker MP, Kennedy MF, Shipley MM, Lindner SE. ALBA4 modulates its stage-specific interactions and specific mRNA fates during Plasmodium yoelii growth and transmission. Mol Microbiol. 2017;106(2):266 –84.

44. Hackmann A, Wu H, Schneider UM, Meyer K, Jung K, Krebber H. Quality control of spliced mRNAs requires the shuttling SR proteins Gbp2 and Hrb1. Nat Commun. 2014;5:3123.

45. Droll D, Wei G, Guo G, Fan Y, Baumgarten S, Zhou Y, et al. Disruption of the RNA exosome reveals the hidden face of the malaria parasite transcriptome. RNA Biol. 2018;15(9):1206 –14.

46. Serpeloni M, Jimenez-Ruiz E, Vidal NM, Kroeber C, Andenmatten N, Lemgruber L, et al. UAP56 is a conserved crucial component of a divergent mRNA export pathway in Toxoplasma gondii. Mol Microbiol. 2016;102(4):672 –89.

47. Simpson CE, Lui J, Kershaw CJ, Sims PF, Ashe MP. mRNA localization to P-bodies in yeast is bi-phasic with many mRNAs captured in a late Bfr1p-dependent wave. Journal of cell science. 2014;127(Pt 6):1254–62.

48. Prommana P, Uthaipibull C, Wongsombat C, Kamchonwongpaisan S, Yuthavong Y, Knuepfer E, et al. Inducible knockdown of Plasmodium gene expression using the glmS ribozyme. PloS one. 2013;8(8):e73783.

49. Labun K, Montague TG, Krause M, Torres Cleuren YN, Tjeldnes H, Valen E. CHOPCHOP v3: expanding the CRISPR web toolbox beyond genome editing. Nucleic acids research. 2019;47(W1):W171–W4.

50. Deitsch K, Driskill C, Wellems T. Transformation of malaria parasites by the spontaneous uptake and expression of DNA from human erythrocytes. Nucleic acids research. 2001;29(3):850 –3.

51. Lambros C, Vanderberg JP. Synchronization of Plasmodium falciparum erythrocytic stages in culture. The Journal of parasitology. 1979;65(3):418 –20.

52. Dobin A. STAR. 2.7.3 ed2019.

53. Andrews S. FastQC. 0.11.8 ed2016.

54. Brennan P. drawProteins: a Bioconductor/R package for reproducible and programmatic generation of protein schematics. F1000Res. 2018;7:1105.

55. Lee S, Zhang A, Su S, Ng A, Holik A, Asselin-Labat M-L, et al. Covering all your bases: incorporating intron signal from RNA-seq data. NAR Genomics and Bioinformatics. 2020;2(3).

56. Katoh K, Standley DM. MAFFT multiple sequence alignment software version 7: improvements in performance and usability. Molecular biology and evolution. 2013;30(4):772 –80.

57. Kearse M, Moir R, Wilson A, Stones -Havas S, Cheung M, Sturrock S, et al. Geneious Basic: an integrated and extendable desktop software platform for the organization and analysis of sequence data. Bioinformatics. 2012;28(12):1647 –9.

58. Capella-Gutierrez S, Silla-Martinez JM, Gabaldon T. trimAl: a tool for automated alignment trimming in large-scale phylogenetic analyses. Bioinformatics. 2009;25(15):1972–3.

59. Stecher G, Tamura K, Kumar S. Molecular Evolutionary Genetics Analysis (MEGA) for macOS. Molecular biology and evolution. 2020;37(4):1237–9.

60. Schindelin J, Arganda-Carreras I, Frise E, Kaynig V, Longair M, Pietzsch T, et al. Fiji: a n open-source platform for biological-image analysis. Nature methods. 2012;9(7):676–82.

